# Kinetics and correlates of the neutralizing antibody response to SARS-CoV-2

**DOI:** 10.1101/2021.01.26.428207

**Authors:** Kanika Vanshylla, Veronica Di Cristanziano, Franziska Kleipass, Felix Dewald, Philipp Schommers, Lutz Gieselmann, Henning Gruell, Maike Schlotz, Meryem S Ercanoglu, Ricarda Stumpf, Petra Mayer, Eva Heger, Wibke Johannis, Carola Horn, Isabelle Suárez, Norma Jung, Susanne Salomon, Kirsten Alexandra Eberhardt, Gerd Fätkenheuer, Nico Pfeifer, Ralf Eggeling, Max Augustin, Clara Lehmann, Florian Klein

## Abstract

A detailed understanding of antibody-based SARS-CoV-2 immunity has critical implications for overcoming the COVID-19 pandemic and for informing on vaccination strategies. In this study, we evaluated the dynamics of the SARS-CoV-2 antibody response in a cohort of 963 recovered individuals over a period of 10 months. Investigating a total of 2,146 samples, we detected an initial SARS-CoV-2 antibody response in 94.4% of individuals, with 82% and 79% exhibiting serum and IgG neutralization, respectively. Approximately 3% of recovered patients demonstrated exceptional SARS-CoV-2 neutralizing activity, defining them as ‘elite neutralizers’. These individuals also possessed effective cross-neutralizing IgG antibodies to SARS-CoV-1 without any known prior exposure to this virus. By applying multivariate statistical modeling, we found that sero-reactivity, age, time since disease onset, and fever are key factors predicting SARS-CoV-2 neutralizing activity in mild courses of COVID-19. Investigating longevity of the antibody response, we detected loss of anti-spike reactivity in 13% of individuals 10 months after infection. Moreover, neutralizing activity had an initial half-life of 6.7 weeks in serum versus 30.8 weeks in purified IgG samples indicating the presence of a more stable and long-term memory IgG B cell repertoire in the majority of individuals recovered from COVID-19. Our results demonstrate a broad spectrum of the initial SARS-CoV-2 neutralizing antibody response depending on clinical characteristics, with antibodies being maintained in the majority of individuals for the first 10 months after mild course of COVID-19.

## Main

COVID-19 is caused by the severe acute respiratory syndrome coronavirus 2 (SARS-CoV-2), which was first identified in December 2019^1,2^. Since then, the virus has rapidly spread across the globe and caused more than 90 million proven infections and over 2 million deaths. Disease severity ranges from asymptomatic infection to symptoms like cough, fever, muscle pain, and diarrhea to severe courses of infection including pneumonia with severe respiratory distress and a high risk of death^3-5^. While the majority of infected individuals experience a mild course of disease, elderly or individuals with pre-existing conditions are at higher risk for severe courses of COVID-19^6^. In symptomatic non-hospitalized cases, the acute course of disease typically spans 7-14 days^7,8^. However, a significant fraction of COVID-19 patients suffer long-lasting symptoms post recovery, so called ‘post-COVID syndrome’^9-11^ (Augustin *et al*., submitted).

SARS-CoV-2 infects human cells by using the virus spike (S) protein^12^ for targeting the angiotensin converting enzyme-2 (ACE-2) receptor^13^. The S-protein carries dominant epitopes against which humoral B and T cell responses are generated upon natural infection and vaccination^14-18^. Spike-specific IgM, IgA, and IgG antibodies are detected early after infection^19,20^ and IgG antibody levels and IgG memory B cells can persist post infection^21^.

Neutralizing antibodies (NAbs) are powerful molecules that target viruses and block infection. Moreover, they can eliminate circulating viruses and infected cells by antibody-mediated effector functions^22,23^. As a result, NAbs are crucial to overcome infectious diseases and are an important correlate of protection^24^. For SARS-CoV-2, vaccine induced NAbs as well as purified IgGs from convalescent animals have been shown to protect non-human primates (NHPs) from infection in a SARS-CoV-2 challenge model^25,26^. Moreover, highly potent monoclonal NAbs have been isolated^27-29^ and are being used for treatment of COVID-19 in humans^30,31^.

Given the short time SARS-CoV-2 has been studied, information on long-term antibody dynamics are limited. Recent studies show that serum neutralizing activity is detectable within a week after onset of symptoms^32,33^ and can persist for the first months after infection^21,23,34^. Moreover, studies with symptomatic and hospitalized individuals have shown that more severe courses of disease result in a stronger SARS-CoV-2 neutralizing antibody response^14,35,36^. While these studies provide important insights, a precise quantification of SARS-CoV-2 neutralizing activity and dynamics as well as clinical correlates of developing a protective antibody response are largely unknown.

In this study, we set out to provide a deeper understanding of the neutralizing antibody response to SARS-CoV-2. To this end, we determined neutralizing serum and IgG activity of 2,146 samples from a longitudinally monitored cohort of 963 individuals over time together with detailed information on the course of disease and past medical history. We combined statistical modeling to infer antibody decay rates after SARS-CoV-2 infection and built a prediction model for evaluating how clinical or disease features correlate with NAb titers. Finally, we performed longitudinal analyses to study anti-spike antibody levels as well as NAb titers for a time period of up to 10 months post SARS-CoV-2 infection. Our results inform on the kinetics, longevity and features affecting the antibody response to SARS-CoV-2. They are critical to understand SARS-CoV-2 immunity and to guide non-pharmacological interventions and vaccination strategies to overcome COVID-19^37^.

## Results

### Establishing a cohort for investigating SARS-CoV-2 immunity

To investigate the development of SARS-CoV-2 immunity, we established a cohort of COVID-19 patients who recently recovered from SARS-CoV-2 infection. Time since disease onset was derived from self-reported symptom onset or date of positive naso-/oro-pharyngeal swab. In addition, each participant reported details on the course of infection, symptoms, and past medical history (**Supplementary Table 1**). Participants enrolled ranged from 18-79 years of age (median: 44 years) with a balanced distribution of males (46.1%) and females (53.9%). Disease severity included asymptomatic (4.6%), mildly symptomatic (91.69%), and hospitalized individuals (2.6%; **Fig. 1, Supplementary Table 1**). 23.4% of participants reported pre-existing conditions that have been described to influence COVID-19 outcomes^6^.

**Figure 1:**
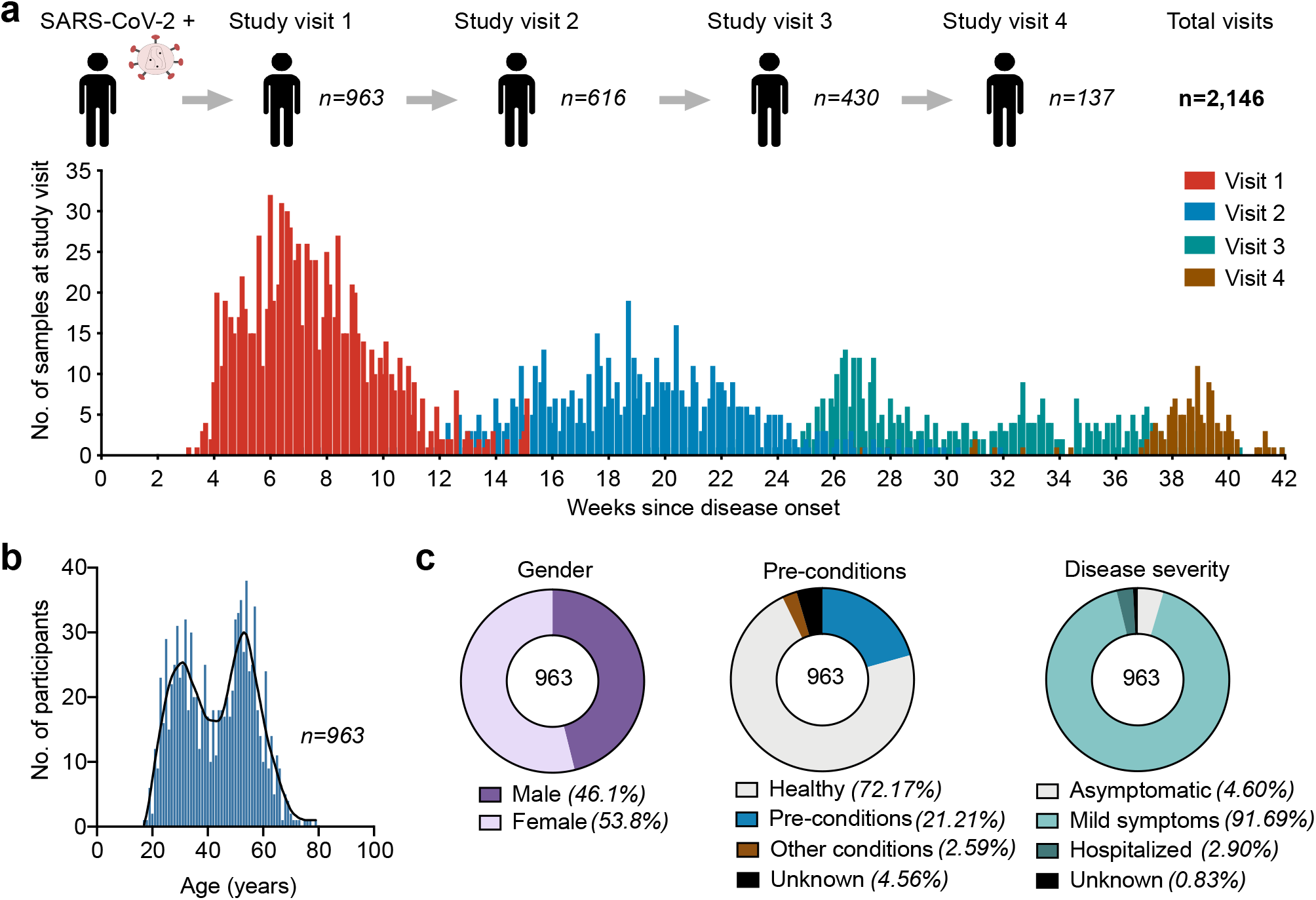
SARS-CoV-2 recovered cohort and study design. **a**, Illustration depicting study timeline and number of individuals analyzed at each study visit. Graph represents sample collection time for participants in weeks since disease onset (symptom onset date or positive PCR date). **b**, age distribution of the cohort **c**, gender distribution, presence of pre-conditions and disease severity.

Blood samples were collected from 963 individuals at study visit 1 (median 7.3 weeks post disease onset) with follow up analyses at study visit 2 for 616 participants (median 18.8 weeks post disease onset), study visit 3 for 430 participants (median 30.1 weeks post disease onset), and study visit 4 for 137 participants (median 37.9 weeks post disease onset; **Fig. 1**). Other participants were lost in follow-up or did not reach the respective study visit at the time of our analysis. Anti-spike IgG was quantified by ELISA and chemiluminescent immunoassays (CLIA) and the NAb response to SARS-CoV-2 was analyzed using both serum dilutions as well as purified IgG to precisely quantify neutralizing activity (**Extended Data Fig. 1**). In total, 4,516 measurements were collected for visit 1 with another 1,867 subsequent measurements for visit 2-4 to determine the SARS-CoV-2 antibody response for 10 months following infection.

### Broad spectrum of the initial SARS-CoV-2 neutralizing antibody response

NAb levels were quantified by testing serum and purified IgG from plasma/serum against pseudovirus particles expressing the Wuhan01 spike protein (EPI_ESL406716). Serum neutralization at study visit 1 was categorized based on titer into non- (ID_50_<10), low- (ID_50_=10-25), average- (ID_50_=25-250), high- (ID_50_=250-2500), and elite-neutralizers (ID_50_>2500; **Fig. 2a**). Mean serum ID_50_ titer was 111.3 with 17.7% of individuals that did not reach 50% neutralization at the lowest serum dilution of 1:10. In addition, all samples were purified for IgG and the neutralizing response was determined and categorized based on IC_50_ values into non- (IC_50_ > 750 μg/ml), low- (IC_50_ = 500-750 μg/ml), average- (IC_50_ = 100-500 μg/ml), high- (IC_50_ = 20-100 μg/ml), and elite-neutralization (IC_50_ < 20 μg/ml; **Fig. 2b**). At study visit 1, out of 963 participants, 10%, 44.8%, and 20% demonstrated low, average, and high neutralization, respectively. 21% did not mount an IgG neutralizing response of an IC_50_ below 750 µg/ml. 3.3% of individuals were classified as ‘elite neutralizers’ with IC_50_ values as low as 0.7 μg/ml detected in one individual at 8.6 weeks post disease onset. Combining serum and IgG measurements, 87.3 % individuals showed detectable NAb activity at median 7.3 weeks after SARS-CoV-2 infection (**Fig. 2c**). The serum and IgG neutralization potency categorization matched for most individuals with a high correlation between serum ID_50_ titers and IgG IC_50_ values (spearman r = -0.72, p < 0.0001; **Fig. 2c**). Moreover, only 5% samples had only serum and no IgG neutralization indicating that IgG antibodies forms the dominant NAb isotype in serum. To further determine the predictive value of IgG binding to the S protein for SARS-CoV-2 neutralization, we performed an S1-reactive ELISA (Euroimmun) on all samples of visit 1. 82.8% and 70.2% of individuals possessed spike-reactive IgG (**Fig. 2d, e)** and IgA Abs, respectively (**Fig. 2d and Extended Data Fig. 2a**). Anti-spike IgG levels were directly proportional to IgG NAb IC_50_ values (spearman r = -0.62, p < 0.0001; **Fig. 2f**) and IgG Ab levels better correlated with serum neutralization than IgA Ab levels (**Extended Data Fig. 2c, d**).

**Figure 2:**
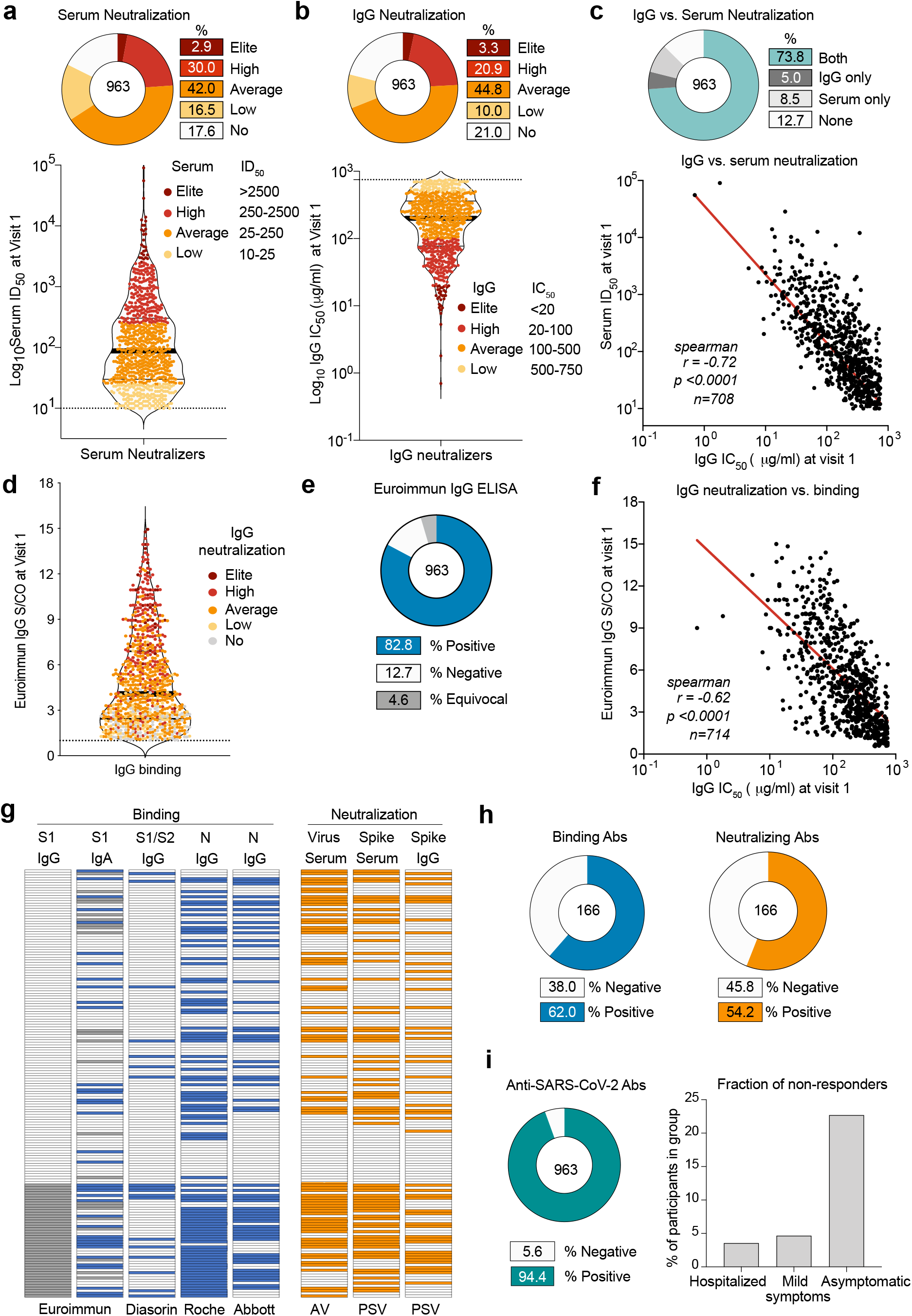
Neutralizing antibody response after recovery from SARS-CoV-2 infection. **a**, pie chart illustrating fraction of serum neutralizers against Wu01 pseudovirus at study visit 1. Violin plot depicts serum ID_50_ values for the neutralizers (*n=793*), categorized based on serum ID_50_ titers. Dotted line represents the LOD (10-fold dilution) of the assay. **b**, pie chart depicting the fraction of IgG neutralization against Wu01 pseudovirus at study visit 1. Violin plot depicts IgG IC_50_ values for the neutralizers (*n=760*), categorized based on IgG IC_50_. Dotted line represents the LOD (750 μg/ml) of the assay. **c**, pie chart comparing fraction of samples with neutralization at serum and/or IgG level. Spearman correlation plot between serum ID_50_ and IgG IC_50_ values at study visit 1. **d**, violin plot of Euroimmun ELISA signal over cut-off (S/CO) ratios for anti-spike IgG. Dotted line represents LOD (S/CO=1.1) of the assay. **e**, pie charts illustrating fraction of anti-spike IgG reactive individuals in the Euroimmun ELISA. **f**, spearman correlation between Euroimmun IgG S/CO and IgG IC_50_ at study visit 1. **g**, plot depicting binding against spike, Nucleocapsid (N) and neutralizing response against authentic virus (AV) and Wu01 pseudovirus (PSV) of the IgG negative fraction (n=166) with each row representing 1 individual. **h**, pie charts showing total fraction of individuals with binding or neutralizing activity in the IgG-fraction from **g. i**, pie chart representing total combined binding and NAb response in the cohort (n=963) and bar graph of the Ab-negative individuals based on disease severity. LOD, limit of detection

Finally, we determined the fraction of individuals lacking any detectable antibody response. To this end, we combined the results of different IgG and IgA assays detecting binding to SARS-CoV-2 S1, S1/S2, and N as well as three neutralization assays (**Fig. 2g**). Out of the 166 anti-S1-IgG negative (12.7%) or equivocal (4.6%) individuals, we found binding antibodies in 62.0% in at least one of four assays and neutralizing activity in 54.2% in at least one of three assays (**Fig. 2g, h**). Combining these results and accounting for assay-specificity (see methods) we show that only 5.6%-7.3% of individuals do not mount a detectable antibody response against SARS-CoV-2. Notably, while only 3.6% (1 of 28) of hospitalized patients and 4.9% (43 of 877) of individuals with mild symptoms lacked anti-SARS-CoV-2 antibodies, 22.7% (10 of 44) asymptomatic individuals were negative for a detectable antibody response in at visit 1. We conclude that 92.7-94.4% of individuals naturally infected with SARS-CoV-2 mount an antibody response against the virus within the first 12 weeks. Among those, we detected a broad variation in neutralizing activity with approximately 3% generating a highly potent serum and IgG NAb response.

### Sero-reactivity, age, and disease severity predict SARS-CoV-2 neutralization

Next, we analyzed how age, disease severity, gender, and the presence of pre-existing conditions correlate with the anti-spike antibody and SARS-CoV-2 neutralizing response (**Fig. 3a, Extended Data Fig. 3**). The IgG NAb response was significantly higher in older individuals (p <0.0001), with participants >60 years comprising 7.7% of elite- and 42.8% of high-neutralizers (**Fig. 3a**). Hospitalized patients and individuals with symptoms had significantly higher NAb activity (p = 0.0008 and 0.0003) compared to asymptomatic individuals, of which 43.2% (25 of 44) lacked detectable IgG NAbs (**Fig. 3a**). Males showed higher SARS-CoV-2 neutralization than females (GeoMean IC_50_ 136.3 μg/ml vs. 188.4 μg/ml; p <0.0001). In addition, individuals with pre-existing conditions had slightly higher NAb activity compared to those without them (GeoMean IC_50_ 161.9 μg/ml vs. 174.6 μg/ml; p = 0.022; **Fig. 3a**). Similar to IgG NAb activity, serum neutralizing activity and anti-spike antibodies were also higher in older individuals, patients with a more severe course of disease, and males (**Extended Data Fig. 3a-c**). Next, we performed a multivariate statistical analysis to determine the interplay between clinical features and the NAb response. Features included gender, age, disease severity, presence of pre-existing conditions, disease symptoms (**Supplementary Table 1**), weeks since disease onset, and the anti-spike IgG/IgA ELISA measurements. We applied a stepwise regression that adds new features only if they significantly improved the model according to a likelihood ratio test. The resulting IC_50_ prediction model (Adjusted R^2^ = 0.461) revealed that IgG antibody levels are most predictive for SARS-CoV-2 neutralizing activity (p = 10^−99^), followed by age (p = 6.1*10^−7^), IgA antibody levels (p = 7.6*10^−6^), time since disease onset (p = 0.01) and fever during infection (p = 0.02; **Fig. 3b, c**). Similarly, age, anti-spike antibody levels, times since disease onset and fever during acute infection were also found to be highly predictive of serum ID_50_ (**Extended Data Fig. 4a, 4b**). Additionally, we built a Bayesian network model to determine the feature dependencies and how they predict the SARS-CoV-2 IgG neutralizing response (**Fig. 3d)**. When applying the stepwise regression model only for predicting the presence of anti-spike antibodies, we observed that gender (IgG p = 8.5*10^−5^; IgA p = 2.2*10^−10^) and the disease symptoms, cough (IgA p = 0.01), diarrhea (IgG p = 0.02) or change in taste (IgG p = 0.002; IgA p = 0.04) are predictive of anti-spike antibody levels (**Extended Data Fig. 4b, c**). In addition, we investigated the possible effect of viral load obtained from naso-/oro-pharyngeal swabs at the time of diagnosis on the antibody response at study visit 1, but no correlation was found (**Extended Data Fig. 4d, e)**. In summary, higher IgG levels, older age and fever during acute infection are highly predictive of the development of SARS-CoV-2 neutralizing activity.

**Figure 3:**
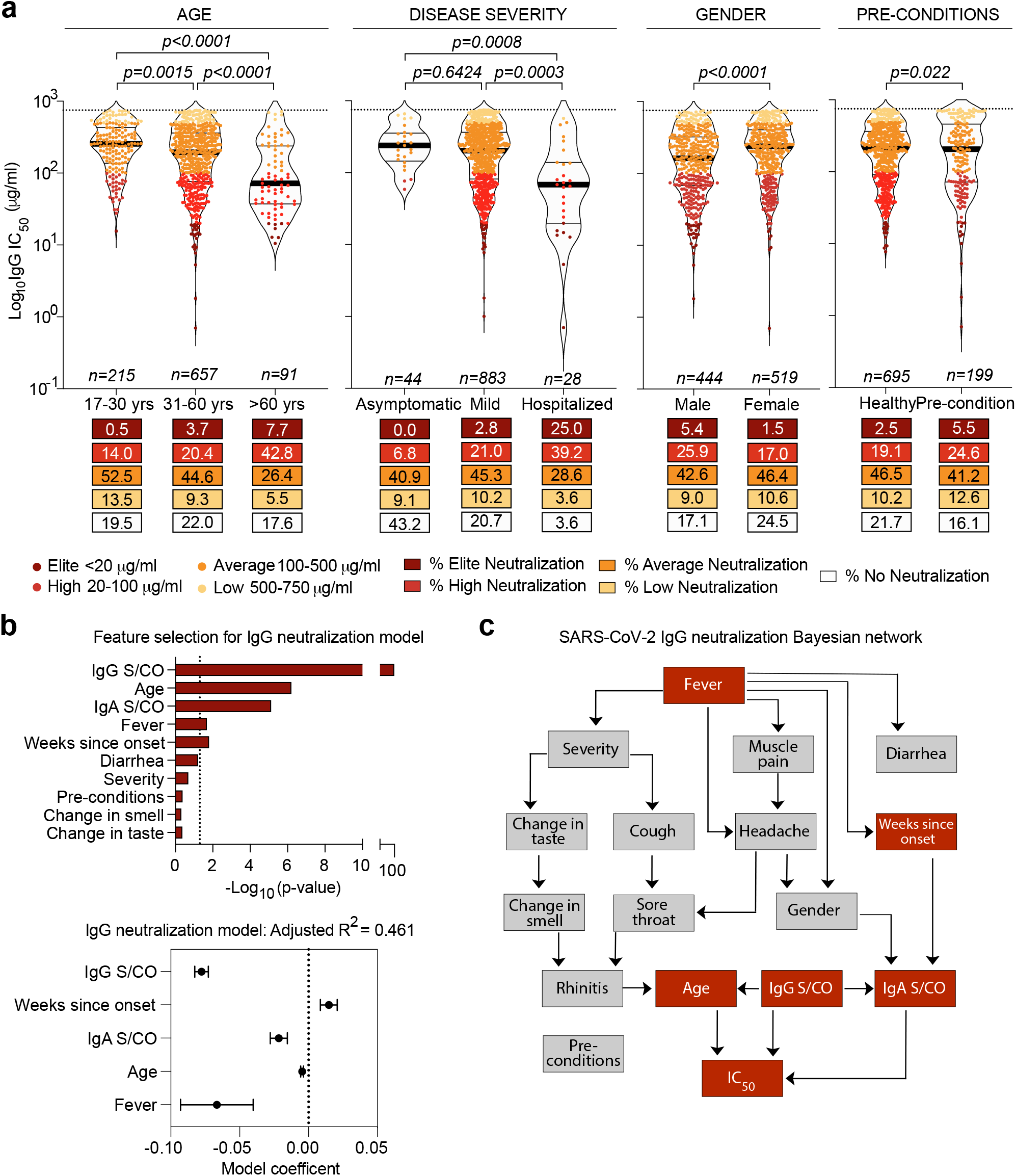
Correlates of neutralizing activity against SARS-CoV-2. **a**, violin plots depicting IgG neutralization IC_50_ values at study visit 1 against Wu01 pseudovirus, subdivided based on age, disease severity, gender and pre-conditions. Dotted line represents the limit of detection (750 μg/ml) of the assay. Statistical analysis was performed Kruskal-Wallis and Mann-Whitney tests. **b**, multiple linear regression model for predicting IgG IC_50_ using the features: Euroimmun S/CO, gender, age, disease severity, pre-conditions, weeks since infection and the 9 reported symptoms. Plot below depicts model coefficients to study the goodness of fit of the final IC_50_ prediction model. **c**, Bayesian network of the features predicting IgG IC_50_ are plotted using the bnlearn R package. The graph connects the features which are predictive of each other with IgG IC_50_ as sink.

### Elite SARS-CoV-2 neutralizers exhibit SARS-CoV-1 cross-neutralization

Individuals mounting a highly potent neutralizing antibody response are often considered ‘elite neutralizers’^38^. These individuals are of particular interest i.) to identify factors associated with the development of effective humoral immunity, ii.) to guide vaccine design, and iii.) to isolate highly potent neutralizing monoclonal antibodies^39^. In order to characterize the small fraction of SARS-CoV-2 elite neutralizers in our cohort (3%; IC_50_ < 20 μg/ml; **Fig. 2b**), we selected 15 individuals of each group of non, low, average, high and elite-neutralizers (**Extended Data Fig. 5a-c**) testing them against authentic SARS-CoV-2 as well as SARS-CoV-1 pseudovirus. Neutralization of SARS-CoV-2 pseudovirus against authentic virus correlated closely in all groups with authentic virus (spearman r = 0.79; **Extended data Fig. 5d**). SARS-CoV-1 neutralization was not observed in non- and low-neutralizers and only in 1 out of 15 average neutralizers. However, in the high and elite neutralizing groups, 8/15 and 15/15 samples carried SARS-CoV-1 cross-neutralizing activity, respectively, with potencies (IC_50_) as low as 5.1 µg/ml IgG. Of note, while all SARS-CoV-2 elite neutralizers demonstrated SARS-CoV-1 cross-neutralization, variation in potency is ranging from 12.1 – 634.9 µg/ml and an overall low correlation (spearman r = 0.3745; **Fig. 4b**). Next, we studied the neutralizing potency of the elite neutralizers against six different SARS-CoV-2 strains carrying several mutations that became prominent at a global level^40^ (**Fig. 4c, Extended Data Fig. 5**). IgG from elite neutralizers was potent against all tested SARS-CoV-2 strains including both S1 and S2 mutants as well as variants (BAVP1, DRC94) carrying the D614G mutation (**Fig. 4c, Extended Data Fig. 5**). We conclude, that individuals mounting a potent SARS-CoV-2 NAb response possess cross-reactive antibodies against SARS-CoV-1 without any known prior exposure and are effective in neutralizing various prevalent SARS-CoV-2 strains.

**Figure 4:**
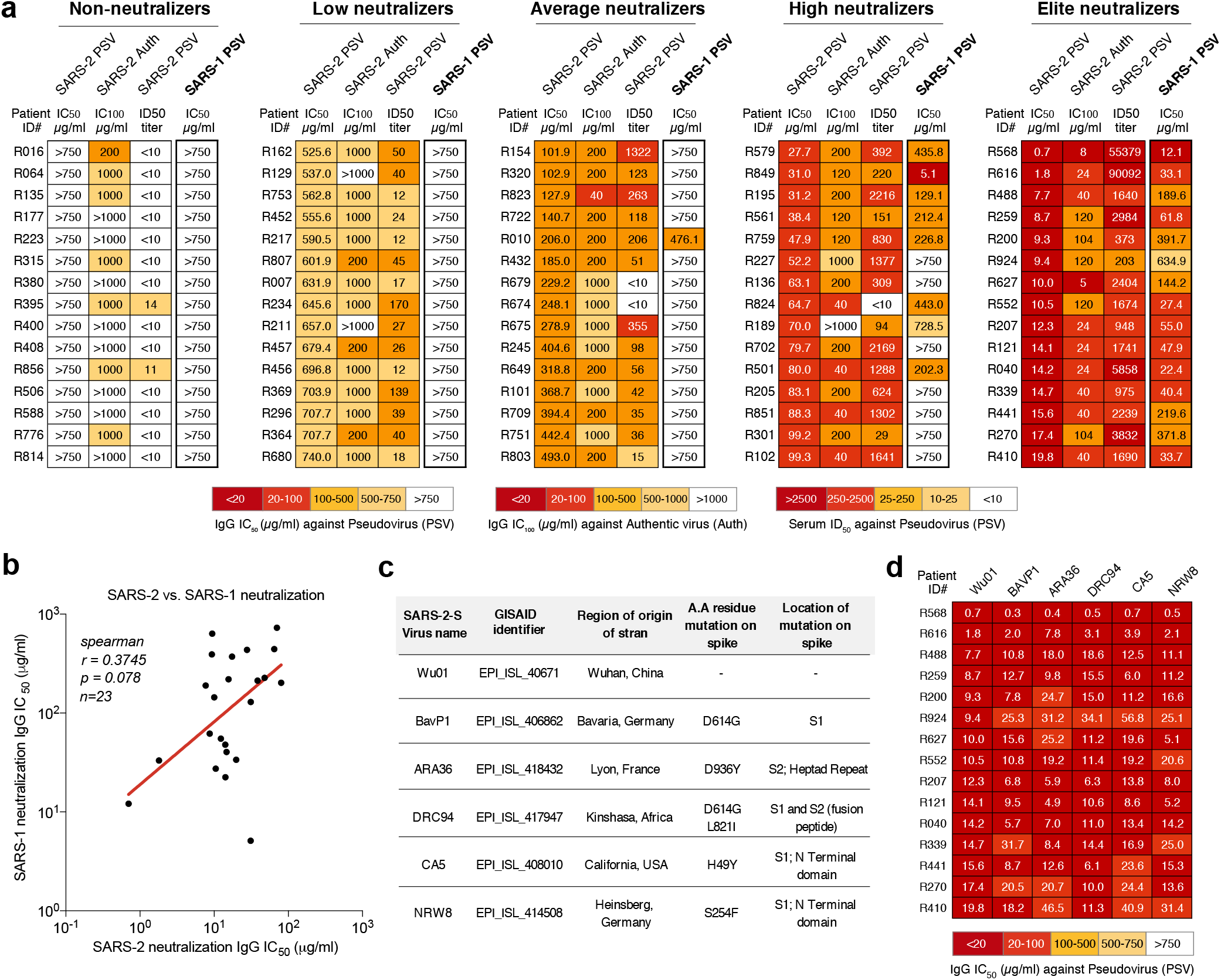
Cross-neutralization by SARS-CoV-2 elite neutralizers. **a**, heat maps visualizing the neutralizing activity of 15 individuals from each neutralization category: Elite-, High-, Average-, Low-, and Non-neutralizers (total n=75) against SARS-CoV-2-S pseudovirus, SARS-CoV-2 authentic virus and SARS-CoV (SARS-1) pseudovirus. **b**, Spearman correlation of IgG IC_50_ against SARS-2-S and SARS-1-S pseudovirus. **c**, details on the source and type of spike mutations in 6 global strains of SARS-CoV-2 generated and used in this study. **d**, heat map visualizing the IC_50_ values of 15 Elite-neutralizers against the 6 SARS-CoV-2 global spike variants from **c**.

### Long-term persistence of IgG NAbs after SARS-CoV-2 infection

In order to study antibody kinetics, we first investigated the development of SARS-CoV-2-directed antibodies in the first 4 weeks after disease onset. To this end, we evaluated 259 samples obtained from an additional 110 individuals. In this subgroup, 44.5% and 54,5% were male and female, respectively, and 41.8% had been hospitalized (**Extended Data Fig. 6a**). Anti-spike IgG and IgA could be detected in some people within the first week after disease onset, with IgA levels starting to decline by week 4 (**Extended Data Fig. 6b**). Out of the 24 individuals that were closely monitored, most individuals sero-converted between 2-3 weeks post disease onset (**Extended Data Fig. 6b**).

In order to assess longevity of humoral immunity following SARS-CoV-2 infection, we applied a linear regression mixed effects model to antibody measurements obtained between 3.1 to 41.9 weeks post infection. The half-life of anti-spike IgG was estimated to be 34.9 weeks (**Fig. 5a**). For systematic tracking of the antibody response within individuals, we analyzed anti-spike antibodies in 131 individuals at 4 study visits (range 3.1 to 38.7 weeks post disease onset; **Fig. 5b, c**). The data show that IgG levels decrease between 1^st^ to 2^nd^ study visit (Geo. Mean S/C0=4.6 vs. Geo. Mean S/C0=3.7) followed by a relatively constant IgG levels for 10 months after infection (Geo. Mean S/C0=3.0) (**Fig. 5b, Supplementary Table 1**). While the detection of S1-reactivity stays equal at first and second visit (86%), the fraction of individuals that are reactive for S1-reactive antibodies decays to 79% (7% drop from visit 1) at the third visit and to 73% (13% drop from visit 1) at visit 4 (9-10 months post disease onset).

**Figure 5:**
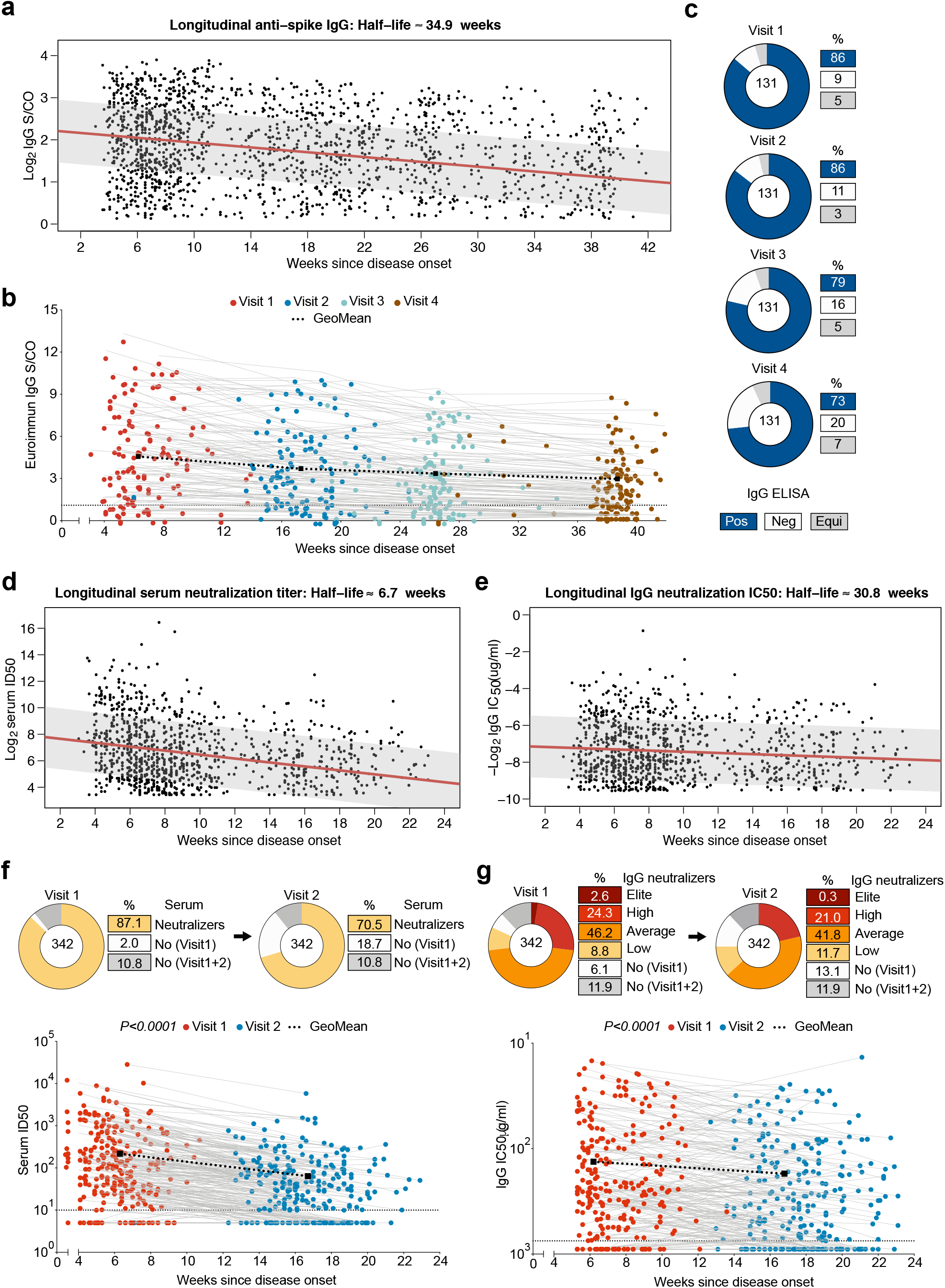
Longitudinal maintenance of anti-SARS-CoV-2 IgG antibody titers. **a**, IgG ELISA ratios (n=1,669) plotted against weeks since infection for half-life estimate of anti-spike IgG levels using a linear mixed-effects model. **b**, longitudinal mapping of IgG levels in 131 individuals from visit 1-4. Dot plots illustrate antibody titer against the weeks since infection to study visit 1 (red) and study visit 2 (blue). Geometric mean change shown in black. Dotted lines represent limit of detection (S/CO=1.1 for IgG ELISA). **c**, pie charts illustrate the change in the fraction of IgG ELISA positive (Pos), Negative (Neg) and Equivocal (Equi) samples (n=131) between the study visits. **d**, serum ID_50_ values against Wu01 pseudovirus (n=1,017) and **e**, IgG IC_50_ values against Wu01 pseudovirus (n=996) plotted against weeks since infection for half-life estimate of the antibody levels using a linear mixed effects model. Longitudinal mapping of serum neutralization (**f**) and IgG neutralization (**g**) in 342 individuals at study visit 1 and 2. Serum and IgG non-neutralizers were assigned values of ID_50_=5 and IC_50_=900 for plotting. Dotted lines represent limit of detection (ID_50_ of 10 and IC_50_ of 750 μg/ml for serum and IgG neutralization assays). Pie charts illustrate the change in the fraction of serum neutralizers (**f**) and IgG neutralizers (**g**) in the samples (n=342) between the study visits.

NAb activity was longitudinally monitored for 342 individuals from visit 1 (median 6.4 weeks post infection) to visit 2 (median 17.3 weeks post infection) (**Fig. 5d-g**). Regression modeling showed that serum NAb titers had a short half-life of 6.7 weeks compared to a much longer 30.8-week half-life for IgG NAb titers (**Fig. 5d, e**). Out of 342 individuals, 87.1% had serum NAb activity at visit 1 whereas only 70.5% had NAb activity remaining at visit 2 (**Fig. 5f**). The overall fraction of IgG neutralizers changed from 82% to 75% between visit 1 and 2. The most dramatic drop from Geo Mean IC_50_ of 16.23 μg/ml to 45.54 μg/ml was detected in elite neutralizers, 88% of whom lost their ‘elite’ status. 23.9% of average/low neutralizers at visit 1 became negative at visit 2 (**Fig. 5g**). Approximately 11% of individuals did not develop any NAbs and remained serum and IgG-negative at both visits. Overall, only 2.4% of the cohort lost detectable antibody responses against SARS-CoV-2 between 1.5 and 4.5 months post infection (**Extended data figure 7a-e)**.

In summary, these results show that in most individuals anti-spike IgG antibody levels are maintained for 10 months with a half-life estimate of 8.7 months. Moreover, even though there is a rapid decline in serum NAb activity, IgG NAb function remains relatively constant with an estimated half-life of 7.7 months. We conclude that although there is a decay of antibody titers in serum, the humoral IgG response persists for as long as 10 months after SARS-CoV-2 infection.

## Discussion

In order to end the COVID-19 pandemic, widespread SARS-CoV-2 protective immunity will be required. Antibodies are critical for effective clearance of pathogens and for prevention of viral infections^41^. In this study, we examined the neutralizing antibody response in 963 individuals who had recently recovered from SARS-CoV-2 infection. The cohort consists primarily (91.69%) of patients with mild COVID-19 therefore representing the predominant clinical course of this disease^6^. Since higher disease severity was shown to correlate with higher antibody responses^14,42^, cohorts mainly composed of hospitalized individuals have limited applicability on the majority of COVID-19 cases^20,35,43,44^. Moreover, to our knowledge this is the most comprehensive study (n=963), in which neutralizing antibody activity has been reported to date with the next largest study having analyzed 4-5 fold less individuals at a single time point^45^.

Upon recovery from COVID-19, we detected the development of a broad spectrum of IgG neutralizing activity ranging from no neutralization (threshold IC_50_ < 750 μg/ml, 21%) to low (IC_50_ = 50-750 μg/ml, 10%), average (IC_50_ = 100-500 μg/ml, 44.8%), high (IC_50_ = 20-100 μg/ml, 20.9%), and elite SARS-CoV-2 neutralization (IC_50_ < 20 μg/ml, 3.3%). 94.4% of individuals were found to possess S- or N-reactive antibodies or neutralizing activity at serum or IgG level. Thus, while most individuals develop a detectable antibody response upon natural infection, the extent of SARS-CoV-2 neutralizing activity is highly variable with the fraction of non-responders being highest for asymptomatic individuals (23%).

The broad spectrum of neutralizing activity developed in COVID-19 recovered individuals may impact the level of protective immunity. For instance, asymptomatic infection is estimated to account for up to 40% of all infections^46^. In these individuals and in other patients with weak antibody responses, lower IgG titers may contribute to a higher susceptibility to re-infection. Recently, mutated virus strains were reported^47,48^, some of which possess mutations causing partial resistance to convalescent plasma^48^ or SARS-CoV-2 monoclonal antibodies^49^. A weak antibody response may help propagate escape variants and may therefore complicate effective measures to combat the COVID-19 pandemic.

To guide vaccine strategies based on population demographics, it is critical to understand which clinical features affect the development of antibody responses. NAb response presented here is comparable to recent spike-based mRNA vaccine studies in age group 18-55, where geometric mean neutralizing titers were in the range of 100-300 ID_50_ (depending on dose) 1.5 months post vaccination^17,50^ versus 111.3 ID_50_ in this study. Recent studies have reported that age, gender and disease severity^14,36,44^ can impact SARS-CoV-2 NAb titers^14,36,42,43,45^. However, a comprehensive analysis on a large representative cohort was missing. Using multivariate statistical analysis on the antibody measurements and clinical data, we found that higher anti-spike antibody levels, older age, symptomatic infection and a severe course of COVID-19 were highly predictive of NAb titers. Notably, based on previous vaccine studies^51^, it was frequently speculated that older individuals might generate a less efficient immune responses to SARS-CoV-2 infection or vaccination. However, based on our data, the >60 age group had the highest level of neutralizing IgG antibodies (mean IC_50_ = 84.8 μg/ml, mean ID_50_ serum titer = 276.6).

In some individuals we detected very high levels of SARS-CoV-2 neutralizing activity (IC_50_ < 20 μg/ml, ID_50_ serum titer > 2,500) ranking them as ‘elite neutralizers’. While cross-reactivity against SARS-CoV-1 and other Beta-CoVs has been shown for some SARS-CoV-2 recovered individuals^52-54^, we revealed that all elite neutralizers have cross-reactive IgG NAbs against SARS-CoV-1. Moreover, IgG from elite neutralizers could efficiently block infection of 6 SARS-CoV-2 strains. Two of them (BavP1 and DRC94) contain the D614G mutation in the S protein^55^ associated with higher infectivity^56^. Given the eminent risk of novel emerging CoVs and monoclonal antibody-resistant SARS-CoV-2 variants, developing antibodies with broader neutralization breadth would be critical. Further evaluation of the antibody response in such elite neutralizers at the single B cell level will be required to understand the details of such potent NAb responses and can yield the identification of new highly potent cross-reactive monoclonal antibodies.

Effective neutralization and clearance of pathogens is mainly mediated by IgG antibodies, which are typically formed within 1-3 weeks post infection and often provide long-term immunity that can last decades^57^. Protective immunity to seasonal coronaviruses like NL63, 229E, OC43 and HKU1 is known to be short lived and re-infection is common^58^. In addition, the antibody response to SARS-CoV-1 and Middle East Respiratory Syndrome (MERS)-CoV was shown to wane over time^59^. Upon SARS-CoV-1 infection, serum IgG and NAbs were shown to decline 3 years after infection^60^. This suggests that immunity to CoVs is rather short lived compared to some other viruses such as measles virus, for which life-long antibody immunity is observed^57^. In our study we not only measured serum neutralization, but also quantified SARS-CoV-2 IgG neutralizing activity. While serum neutralization waned quickly (half-life of 1.5 months), levels of purified IgG rather persisted with a longer half-life of 7.7 months. The sharp drop in serum neutralization could be a consequence of a decline in anti-spike IgA and IgM titers^34^, which along with IgG, cumulatively contribute to serum NAb activity^61^. Finally, SARS-CoV-2 spike-based mRNA vaccines^17^ were shown to induce NAb titers in different age groups for a time span up to 4.25 months^18^. In this study, we found that although SARS-CoV-2-reactive IgG levels decline by 17% within the first 4 months after infection, anti-spike IgG can be persistently detected in the majority of COVID-19 cases for up to 10 months post infection.

In summary, the data presented in this study provides new insight into the features that shape the SARS-CoV-2 NAb response in COVID-19 recovered individuals. Longitudinal mapping of antibody responses reveals a relatively long-lived IgG antibody response lasting up to 10 months. Since many SARS-CoV-2 vaccines are spike protein-based^62^, studying antibody dynamics informs us on longevity of natural immunity as well as may help to inform on vaccination strategies and outcomes in the population.

## Methods

### Enrollment of participants and study design

Blood samples were collected from donors who gave their written consent under the protocols 20-1187 and 16-054, approved by the Institutional Review Board (IRB) of the University Hospital Cologne. All samples were handled according to the safety guidelines of the University Hospital Cologne. Individuals that met the inclusion criteria of i.) ≥18 years old and ii.) history of SARS-CoV-2 positive polymerase chain reaction (PCR) from nasopharyngeal swab or collected sputum, and/or iii.) an onset of COVID-19 specific symptoms longer than 3 weeks ago, were enrolled in this study. Demographical data, COVID-19-related pre-existing conditions, and information on the clinical course were collected at study visit 1. Blood samples were collected starting from study visit 1, for up to 4 follow up visits between the 6^th^ of April and 17^th^ of December 2020.

### Processing of serum, plasma and whole blood samples

Blood samples were collected in Heparin syringes or EDTA monovette tubes (Becton Dickinson) and fractionated into plasma and peripheral blood mononuclear cell (PBMC) by density gradient centrifugation using Histopaque-1077 (Sigma). Plasma aliquots were stored at -80°C till use. Serum was collected from Serum-gel tubes (Sarstedt) by centrifugation and stored at -80°C till use.

### Isolation of IgGs from serum and plasma samples

For the isolation of total IgG, 0.5-1 mL plasma or serum was heat inactivated at 56°C for 45 minutes and incubated overnight with Protein G Sepharose 4 Fast Flow beads (GE Healthcare) at 4°C. Next day, beads were washed on chromatography columns (BioRad) and Protein G bound IgG was eluted using 0.1M Glycine pH=3 and instantly buffered in 1M Tris pH=8. Buffer exchange to PBS (Gibco) was performed using 30 kDa Amicon Ultra-15 columns (Millipore) and the purified IgG was stored at 4°C.

### Cloning of SARS-CoV-2 spike variants

The codon optimized SARS-CoV-2 Wu01 spike (EPI_ISL_40671) was cloned into pCDNA™3.1/V5-HisTOPO vector (Invitrogen). SARS-2-S global strains (BavP1 EPI_ISL_406862; ARA36 EPI_ISL_418432; DRC94 EPI_ISL_417947; CA5 EPI_ISL_408010; NRW8 EPI_ISL_414508) were generated by introducing the corresponding amino acid mutations (**Extended Data Fig. 5**) using the Q5® Site-Directed Mutagenesis Kit (NEB) and per manufacturer’s protocol.

### Production of SARS-CoV pseudovirus particles

Pseudovirus particles were generated by co-transfection of individual plasmids encoding HIV-1 Tat, HIV-1 Gag/Pol, HIV-1 Rev, luciferase followed by an IRES and ZsGreen, and the SARS-CoV-2 spike protein as previously described^63^. In brief, HEK 293T cells were transfected with the pseudovirus encoding plasmids using FuGENE 6 Transfection Reagent (Promega). The virus culture supernatant was harvested at 48h and 72h post transfection and stored at -80°C until use. Each virus batch was titrated by infecting 293T-ACE2 and after a 48-hour incubation period at 37°C and 5% CO_2_, luciferase activity was determined after addition of luciferin/lysis buffer (10 mM MgCl2, 0.3 mM ATP, 0.5 mM Coenzyme A, 17 mM IGEPAL (all Sigma-Aldrich), and 1 mM D-Luciferin (GoldBio) in Tris-HCL) using a microplate reader (Berthold). An RLU of approximately 1000-fold in infected cells versus non-infected cells was used for neutralization assays.

### Pseudovirus assay to determine IgG/plasma/serum SARS-CoV-2 neutralizing activity

For testing SARS-CoV-2 neutralizing activity of IgG or serum/plasma samples, serial dilutions of IgG or serum/plasma (heat inactivated at 56°C for 45 min) were co-incubated with pseudovirus supernatants for 1 h at 37°C prior to addition of 293T cells engineered to express ACE2^63^. Following a 48-hour incubation at 37°C and 5% CO_2_, luciferase activity was determined using the reagents described above. After subtracting background relative luminescence units (RLUs) of non-infected cells, 50% inhibitory concentrations (IC50s) were determined as the IgG concentrations resulting in a 50% RLU reduction compared to untreated virus control wells. 50% Inhibitory dose (ID_50_) was determined as the serum dilution resulting in a 50% reduction in RLU compared to the untreated virus control wells. Each IgG and serum sample were measured in two independent experiments on different days and the average IC_50_ or ID_50_ values have been reported. For each run, a SARS-CoV-2 neutralizing monoclonal antibody was used as control to ensure consistent reproducibility in experiments carried out on different days. Assay specificity calculated using pre-COVID-19 samples was found to be 100%. IC_50_ and ID_50_ values were calculated in GraphPad Prism 7.0 by plotting a dose response curve.

### SARS-CoV-2 live virus isolation from nasopharyngeal swabs

For outgrowth cultures of authentic SARS-CoV-2 from nasopharyngeal swabs, 1×10^6^ VeroE6 cells were seeded onto a T25 flask (Sarstedt) on the previous day DMEM (Gibco) containing 10% FBS, 1% PS, 1mM L-Glutamine and 1mM Sodium pyruvate. 0.2 mL swab in VNT medium was diluted with 0.8 mL DMEM (Gibco) containing 2% FBS, 1% PS, 1mM L-Glutamine and 1mM Sodium pyruvate. The swab dilution was added to VeroE6 cells and left for 1 hour at 37°C, 5%CO_2_ after which an additional 3 mL medium was added. The cultures were examined for the next days for CPE and samples were sent for viral load analysis to track growth of virus by E-gene qPCR. Cell culture supernatant was harvested from positive cultures and stored at -150°C until use. Virus was titrated by adding serial dilutions of virus supernatant (8 replicates) on VeroE6 cells in DMEM (Gibco) containing 2% FBS, 1% PS, 1mM L-Glutamine and 1mM Sodium pyruvate. After 4 days of incubation at 37°C, 5% CO_2_, the presence or absence of cytopathic effects (CPE) was noted in using a brightfield microscope. TCID_50_ was calculated using the Spearman and Kaerber algorithm^64,65^.

### SARS-CoV-2 live virus neutralization assay

Live SARS-CoV-2 (termed CoV2-P3) was grown out from a swab from Cologne using VeroE6 cells as described above and then expanded in culture by superinfection of VeroE6 from the initial outgrowth culture. Whole genome sequencing of the isolated virus was done isolating viral RNA using the QIAamp MinElute Virus Spine kit (Qiagen) and performing Illumina sequencing. The virus spike amino acid sequence (**Extended Data Fig. 5**) is similar to the Wu01 spike (EPI_ISL_40671) with the exception that it contains the D641G mutation. For the neutralization assay, dilutions of IgG were co-incubated with the virus (1000-2000 TCID_50_) for 1 h at 37°C prior to addition of VeroE6 cells in DMEM (Gibco) containing 2% FBS, 1% PS, 1mM L-Glutamine and 1mM Sodium pyruvate. After 4 days of incubation at 37°C, 5% CO_2_, neutralization was analyzed by observing cytopathic effects (CPE) using a brightfield microscope and the highest dilution well with no CPE was noted to be the IC_100_ for the antibody. Assay specificity calculated using pre-COVID-19 samples was found to be 100%. All samples were measured in two independent experiments on separate days and the average IC_100_ from all measurements has been reported.

### Detection of anti-SARS-CoV-2 spike IgG and IgA by ELISA

For assessing IgA and IgG antibody titers, the Euroimmun anti-SARS-CoV-2 ELISA using the S1 domain of the spike protein as antigen was used (Euroimmun Diagnostik, Lübeck, Germany). Serum or plasma samples were tested on the automated system Euroimmun Analyzer I according to manufacturer’s recommendations. Signal-to-cut-off (S/CO) ratio was calculated as extinction value of patient sample/extinction value of calibrator. IgA and IgG S/CO values were interpreted as positive S/CO ≥1.1, equivocal S/CO ≥0.8 - <1.1, and negative S/CO <0.8. Additional commercial kits used for antibody measurements were also used as per manufacturer’s recommendations; Anti-S1/S2 IgG was measured using DiaSorin’s LIAISON® SARS-CoV-2 ELISA kit with the following cut-off values: negative <12.0 AU/ml, equivocal ≥12.0- < 15.0 AU/ml and positive ≥15.0 AU/ml. Anti-N Pan-Igs were measured using Roche’s Elecsys®-Assay with cut-off values: non-reactive < 1,0 COI and reactive ≥ 1,0 COI. Anti-N IgG were measured with Abbott’s Alinity i system with cut-off values: positive S/CO ≥1.4 and negative S/CO <1.4. Assay specificities calculated using pre-COVID-19 samples: Euroimmun IgG 100%; Euroimmun IgA 96%; Roche 98%; Diasorin 98%; Abbott 98%.

### Measurement of SARS-CoV-2 RNA levels from nasopharyngeal swabs

Cycle threshold values for quantifying viral load in naso/oro-pharyngeal swabs was done by qPCR using LightMix® SarbecoV E-gene^66^ plus EAV control (TIB Molbiol, Berlin, Germany) in combination with the N-gene (inhouse primer sets in multiplex PCR) on LightCycler® 480 (Roche Diagnostics).

### Statistical modeling

To select features that are predictive for the log_10_ response in a multivariate analysis (Fig. 3b), forward stepwise regression was applied, using the p-value from a likelihood ratio test (R function lmtest::lrtest) as selection criterion in each step. The final multiple linear regression model (Fig. 3c) includes only features that show a significant model improvement (alpha=0.05) in the feature selection phase. To study the interplay of the different features regarding their relationship with the response (Fig. 3d), a Bayesian network was learned by maximizing the BIC score for hybrid networks via hill-climbing (R function bnlearn::hc)^67^. To enforce it to be a sink in the network, all outgoing edges from the response variable were blacklisted prior to learning. For the longitudinal analyses (Fig. 5e-h), linear mixed effect models (R-function nlme:lme) were applied to all data points from both visits, where each patient has its own intercept. Since a binary transformation of the response was used, half-life estimates were computed as negative inverse of the common slope regression coefficient. Prediction intervals were computed using R-function ggeffects::ggpredict^68^.

## Figure legends

**Extended Data Figure 1: Samples used for analysis of SARS-CoV-2 antibody response**

**a**, Illustration depicting processing of blood samples and IgG purification from plasma or serum samples. **b**, Plot analyzing the efficiency of IgG purification from plasma or serum as compared to clinical reference range. Statistical testing performed with Kruskal-Wallis test. Validation of the pseudovirus neutralization test against SARS-2-S Wu01 pseudovirus using Pre-COVID-19 plasma (**c**) and IgG (**d**) samples with a neutralizing monoclonal antibody as positive control^28^.

**Extended Data Figure 2: Correlation between neutralization and serology results**

**a**, violin plot of Euroimmun ELISA signal over cut-off (S/CO) ratios for anti-spike IgA. Dotted line represents the limit of detection (S/CO=1.1) of the assay. **b**, Spearman correlation plot between Euroimmun IgA S/CO and serum ID_50_ values at study visit 1. Euroimmun IgA S/CO and serum ID_50_ values at study visit 1. Pie charts illustrating the fraction of serum neutralizers and non-neutralizers and their corresponding Euroimmun IgA ELISA result for comparison. **c**, Spearman correlation plot of Euroimmun IgG S/CO ratios vs. IgA S/CO ratios at study visit 1. **d**, Spearman correlation plot between Euroimmun IgG S/CO and serum ID_5_0 values at study visit 1. Pie charts illustrating the fraction of serum neutralizers and non-neutralizers and their corresponding Euroimmun IgG ELISA result for comparison.

**Extended Data Figure 3: Correlates of anti-SARS-CoV-2 antibody titers**

**a**, violin plots depicting serum neutralization at study visit 1 against Wu01 pseudovirus, subdivided based on age, disease severity, gender and pre-conditions. Dotted line represents the limit of detection (1:10 dilution) of the assay. **b**, violin plots depicting Euroimmun IgG ELISA S/CO at study visit 1, subdivided based on age, disease severity, gender and pre-conditions. Dotted line represents the limit of detection (S/CO=1.1) of the assay. **c**, violin plots depicting Euroimmun IgA ELISA S/CO at study visit 1, subdivided based on age, disease severity, gender and pre-conditions. Dotted line represents the limit of detection (S/CO=1.1) of the assay. **a, b** and **c** Statistical analysis was performed using Kruskal-Wallis and Mann-Whitney tests.

**Extended Data Figure 4: Statistical predication of SARS-CoV-2 antibody responses**

**a**, multiple linear regression model and model coefficients for predicting serum neutralization using the features: Euroimmun S/CO, gender, age, disease severity, pre-conditions, weeks since infection and the 9 reported symptoms. **b**, Bayesian network of the features predicting serum ID_50_ are plotted using the bnlearn R package. The graph connects the features which are predictive of each other with serum ID_50_ as sink. **c** and **d**, Multiple linear regression model for predicting IgG and IgA ratios using the features: gender, age, disease severity, pre-conditions, weeks since infection and the 9 reported symptoms. Plots on the right depicts model coefficients to study the goodness of fit of the corresponding final models. Spearman correlation plot for diagnostic naso-/oro-pharyngeal swab Ct values for E-gene (**e**) or N-gene (**f**) vs. IgG IC_50_, serum ID_50_, anti-spike IgG and anti-spike IgA values at study visit 1.

**Extended Data Figure 5: Neutralization of different strains by SARS-CoV-2 elite-neutralizers**

**a**, plots for the distribution of gender, age and time since infection for the 15 individuals selected randomly from the five IgG neutralization categories: elite-, high-, average-, low-, and non-neutralizers (n=75 total). Statistical testing performed with Kruskal-Wallis test using Dunn’s multiple comparisons. **b**, Spearman correlation of IgG IC50 against SARS-2-S pseudovirus and SARS-2 authentic virus. **c**, Relative infectivity of SARS-CoV-2 global strain pseudovirus in 293T-ACE2 cells. **d**, Sequence alignment of the spike amino acid sequence of the 6 global SARS-CoV-2 strains and SARS-1 used for pseudovirus neutralization assays in this study.

**Extended Data Figure 6: Antibody kinetics in the early phase of SARS-CoV-2 infection**

**a**, Pie charts indicating distribution of gender and disease severity in individuals who were longitudinally monitored starting from the early phase of infection. **b**, Plots depicting IgG and IgA ratios over time in individuals (n=107). Dotted line represents the limit of detection (S/CO=1.1) of the Euroimmun ELISA. Statistical analysis was performed using a second order polynomial quadratic equation (R^2^=0.128 for IgG and R^2^=0.140 for IgA) with 95% confidence interval shading (IgG in blue and IgA in red) of the best line. **c**, Individual plots depicting IgG (blue) and IgA (red) levels over time. Gender and disease severity are indicated within each plot. Dotted line represents the limit of detection (S/CO=1.1) of the Euroimmun ELISA.

**Extended Data Figure 7: Changes in Ab response against SARS-CoV-2 over time**

**a**, plot depicting SARS-CoV-2 S1 binding and Wu01 pseudovirus neutralization of 339 individuals at visit 1 and visit 2 with each row representing 1 individual. Bar graphs showing change in fraction of individuals negative for anti-spike Abs (**b**), anti-spike NAbs (**c**) or any Ab response (**d**). **e**, pie chart evaluating the total presence if Ab response between visit 1 and 2 for all individuals.

## Data availability statement

All data including virus spike sequences are available in the manuscript main figures or supplementary material.

## Acknowledgements

We are extremely grateful to all study participants who took part in this study; members of the Klein lab for helpful discussions; Reinhild Brinker, Marie Wunsch and Maike Wirtz for technical support; Daniela Weiland and Nadine Henn for project and laboratory management support; Stefan Poehlmann and Markus Hoffmann for sharing the Wuhan SARS-2-S spike construct; Jesse Bloom and Kate Crawford for sharing 293T-ACE2 cells and lentiviral constructs for production of SARS-CoV pseudovirus particles; Jason McLellan and Nianshuang Wang for sharing the SARS-1-S spike construct; Stephan Becker and Verena Kraehliv for sharing VeroE6 cells; Jeorg Timm, Andreas Walker and Max Damagnez for SARS-CoV-2 virus genome sequencing. This work was funded by grants to Florian Klein from the German Center for Infection Research (DZIF), the German Research Foundation (DFG) CRC1279 and CRC1310, European Research Council (ERC) ERC-stG639961 and COVIM: „NaFoUniMedCovid19“ (FKZ: 01KX2021).

## Author contributions

F.Klein and K.V. conceptualized and designed the study; F.Klein, C.L., G.F. and N.P. provided supervision; K.V. and F.Klein wrote the first draft of the manuscript, all authors reviewed the manuscript draft and agree to the final version; M.A., P.S., L.G., F.D., V.C., H.G., C.H., I.S., N.J., were involved in study participant interaction including obtaining informed consent, clinical data and sample collection and writing the study protocol. K.V. and F.Kleipass performed neutralization assays; V.C. and W.J. obtained ELISA data; F.Kleipass and K.V. performed IgG purification; K.V., V.C., F.K., F.D., P.S. analyzed data; K.V. performed final data analysis and R.E. performed statistical analysis; M.S., M.S.E., R.S. and P.M. processed blood samples; K.E., S.S. and E.H. were involved in data collection.

