## Supplement for "Kinetics and correlates of the neutralizing antibody response to SARS-CoV-2"

#### Extended data figures


Extended data figure 1 *Samples used for analysis of SARS-CoV-2 antibody response*

Extended data figure 2 *Correlation between neutralization and serology results*

Extended data figure 3 *Correlates of anti-SARS-CoV-2 antibody titers*

Extended data figure 4 *Statistical predication of SARS-CoV-2 antibody responses*

Extended data figure 5 *Neutralization of different strains by SARS-CoV-2 elite-neutralizers*

Extended data figure 6 *Antibody kinetics in the early phase of SARS-CoV-2 infection*

Extended data figure 7 *Changes in Ab response against SARS-CoV-2 over time*


#### Supplementary Tables


Supplementary table 1 *Study cohort demographics, clinical information and antibody measurements*

### Extended Data Figure 1

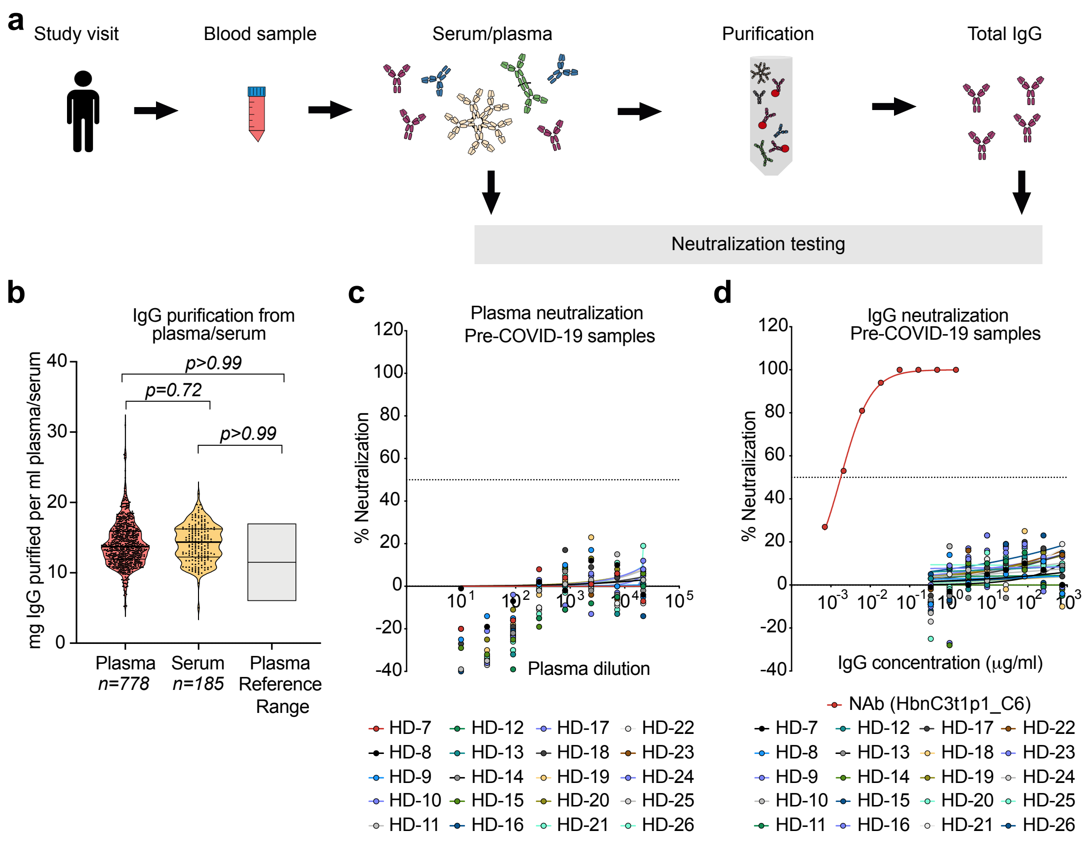


**Extended Data Figure 1: Samples used for analysis of SARS-CoV-2 antibody response**
**a,** Illustration depicting processing of blood samples and IgG purification from plasma or serum samples. **b**, Plot analyzing the efficiency of IgG purification from plasma or serum as compared to clinical reference range. Statistical testing performed with Kruskal-Wallis test. Validation of the pseudovirus neutralization test against SARS-2-S Wu01 pseudovirus using Pre-COVID-19 plasma (**c**) and IgG (**d**) samples with a neutralizing monoclonal antibody as positive control.

### Extended data figure 2

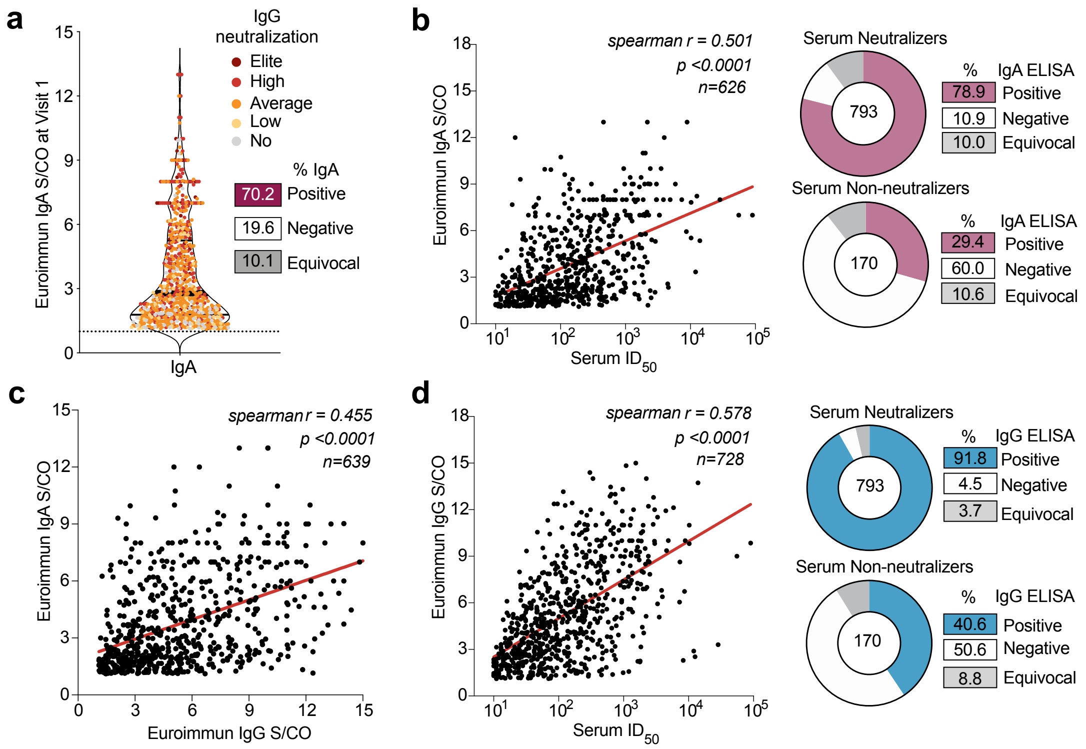


**Extended Data Figure 2: Correlation between neutralization and serology results**
**a,** violin plot of Euroimmun ELISA signal over cut-off (S/CO) ratios for anti-spike IgA. Dotted line represents the limit of detection (S/CO=1.1) of the assay. **b**, Spearman correlation plot between Euroimmun IgA S/CO and serum ID50 values at study visit 1. Euroimmun IgA S/CO and serum ID50 values at study visit 1. Pie charts illustrating the fraction of serum neutralizers and non-neutralizers and their corresponding Euroimmun IgA ELISA result for comparison. **c**, Spearman correlation plot of Euroimmun IgG S/CO ratios vs. IgA S/CO ratios at study visit 1. **d**, Spearman correlation plot between Euroimmun IgG S/CO and serum ID50 values at study visit 1. Pie charts illustrating the fraction of serum neutralizers and non-neutralizers and their corresponding Euroimmun IgG ELISA result for comparison.

### Extended data figure 3

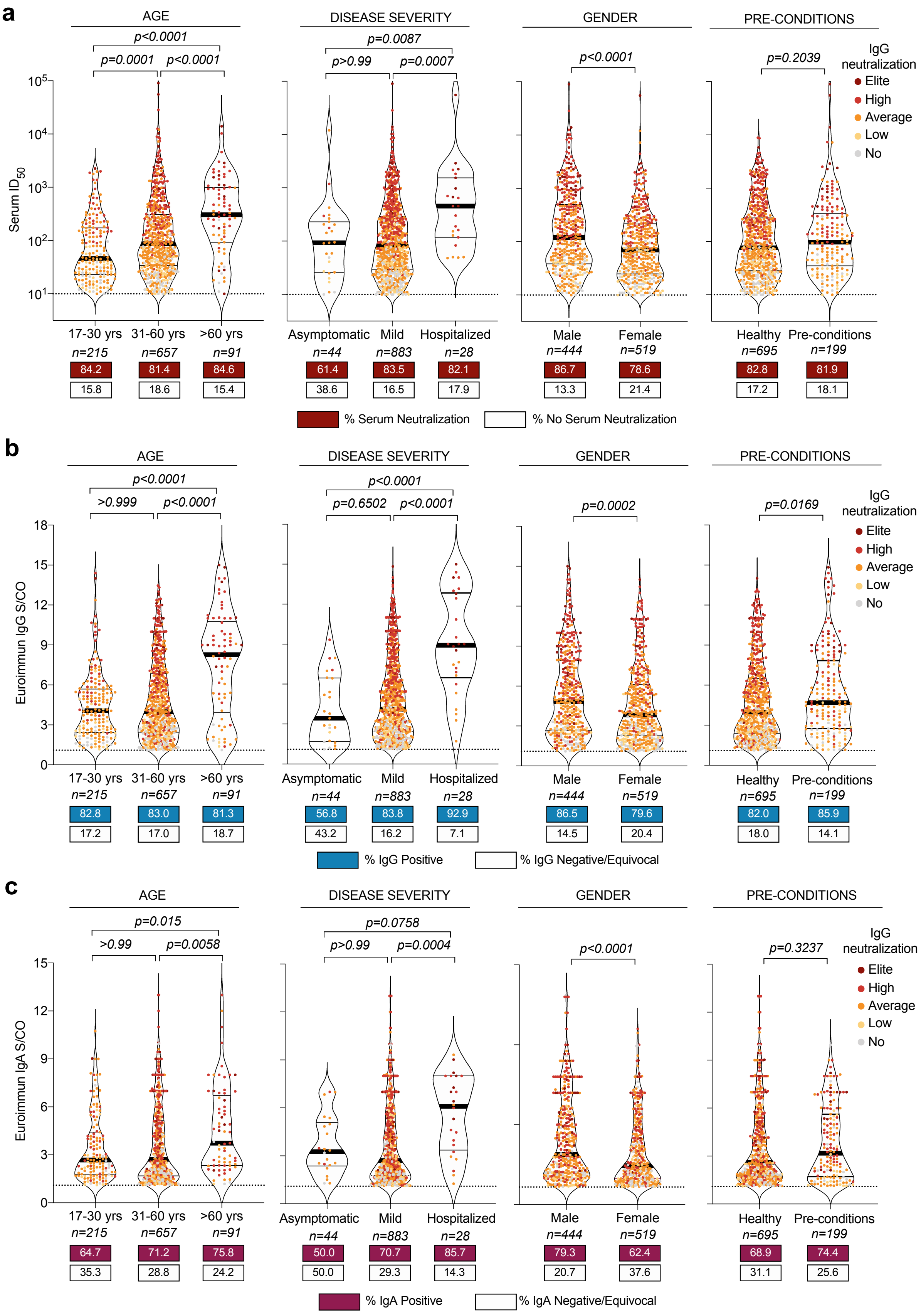


**Extended Data Figure 3: Corelates of anti-SARS-CoV-2 antibody titers**
**a,** violin plots depicting serum neutralization at study visit 1 against Wu01 pseudovirus, subdivided based on age, disease severity, gender and pre-conditions. Dotted line represents the limit of detection (1:10 dilution) of the assay. **b**, violin plots depicting Euroimmun IgG ELISA S/CO at study visit 1, subdivided based on age, disease severity, gender and pre-conditions. Dotted line represents the limit of detection (S/CO=1.1) of the assay. **c**, violin plots depicting Euroimmun IgA ELISA S/CO at study visit 1, subdivided based on age, disease severity, gender and pre-conditions. Dotted line represents the limit of detection (S/CO=1.1) of the assay. **a, b** and **c** Statistical analysis was performed using Kruskal-Wallis and Mann-Whitney tests.

### Extended Data Figure 4

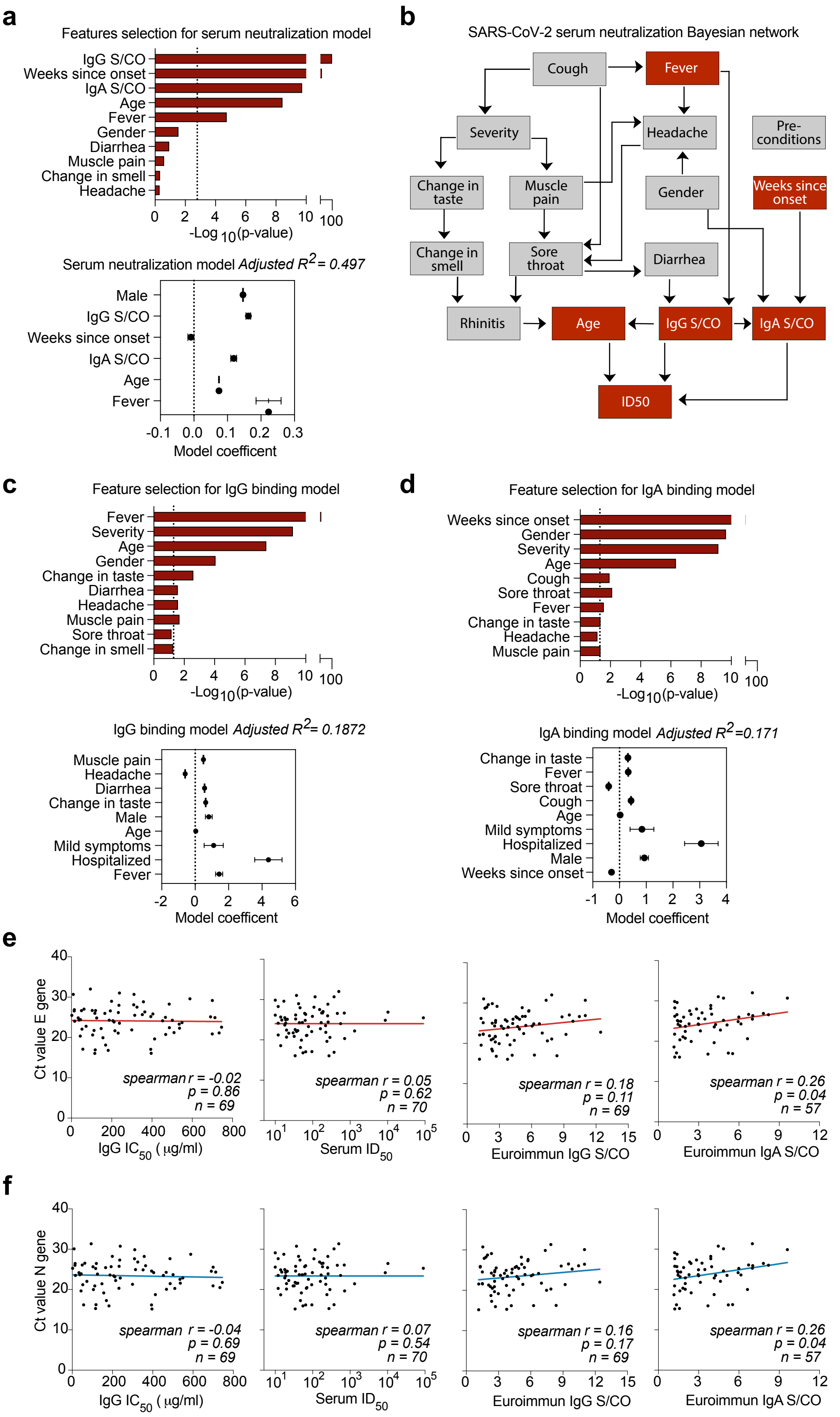


**Extended Data Figure 4: Statistical predication of SARS-CoV-2 antibody responses**
**a,** multiple linear regression model and model coefficients for predicting serum neutralization using the features: Euroimmun S/CO, gender, age, disease severity, pre-conditions, weeks since infection and the 9 reported symptoms. **b**, Bayesian network of the features predicting serum ID50 are plotted using the bnlearn R package. The graph connects the features which are predictive of each other with serum ID50 as sink. **c** and **d**, Multiple linear regression model for predicting IgG and IgA ratios using the features: gender, age, disease severity, pre-conditions, weeks since infection and the 9 reported symptoms. Plots on the right depicts model coefficients to study the goodness of fit of the corresponding final models. Spearman correlation plot for diagnostic naso-/oro-pharyngeal swab Ct values for E-gene (**e**) or N-gene (**f**) vs. IgG IC50, serum ID50, anti-spike IgG and anti-spike IgA values at study visit 1.

### Extended Data Figure 5

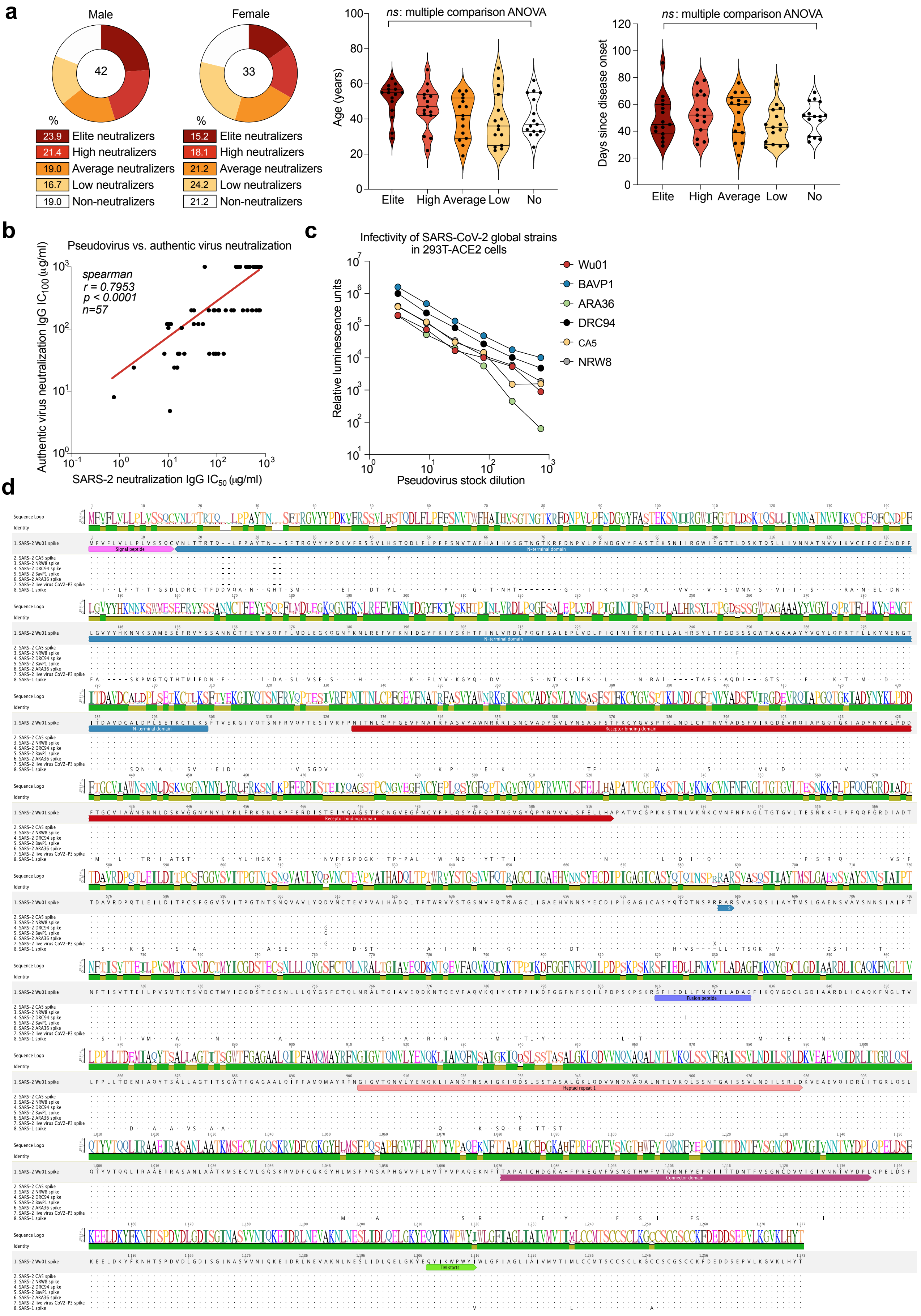


**Extended Data Figure 5: Neutralization of different strains by SARS-CoV-2 elite-neutralizers**
**a,** plots for the distribution of gender, age and time since infection for the 15 individuals selected randomly from the five IgG neutralization categories: elite-, high-, average-, low-, and non-neutralizers (n=75 total). Statistical testing performed with Kruskal-Wallis test using Dunn's multiple comparisons. ***b***, Spearman correlation of IgG IC50 against SARS-2-S pseudovirus and SARS-2 authentic virus. **c**, Relative infectivity of SARS-CoV-2 global strain pseudovirus in 293T-ACE2 cells. **d**, Sequence alignment of the spike amino acid sequence of the 6 global SARS-CoV-2 strains and SARS-1 used for pseudovirus neutralization assays in this study.

### Extended data figure 6

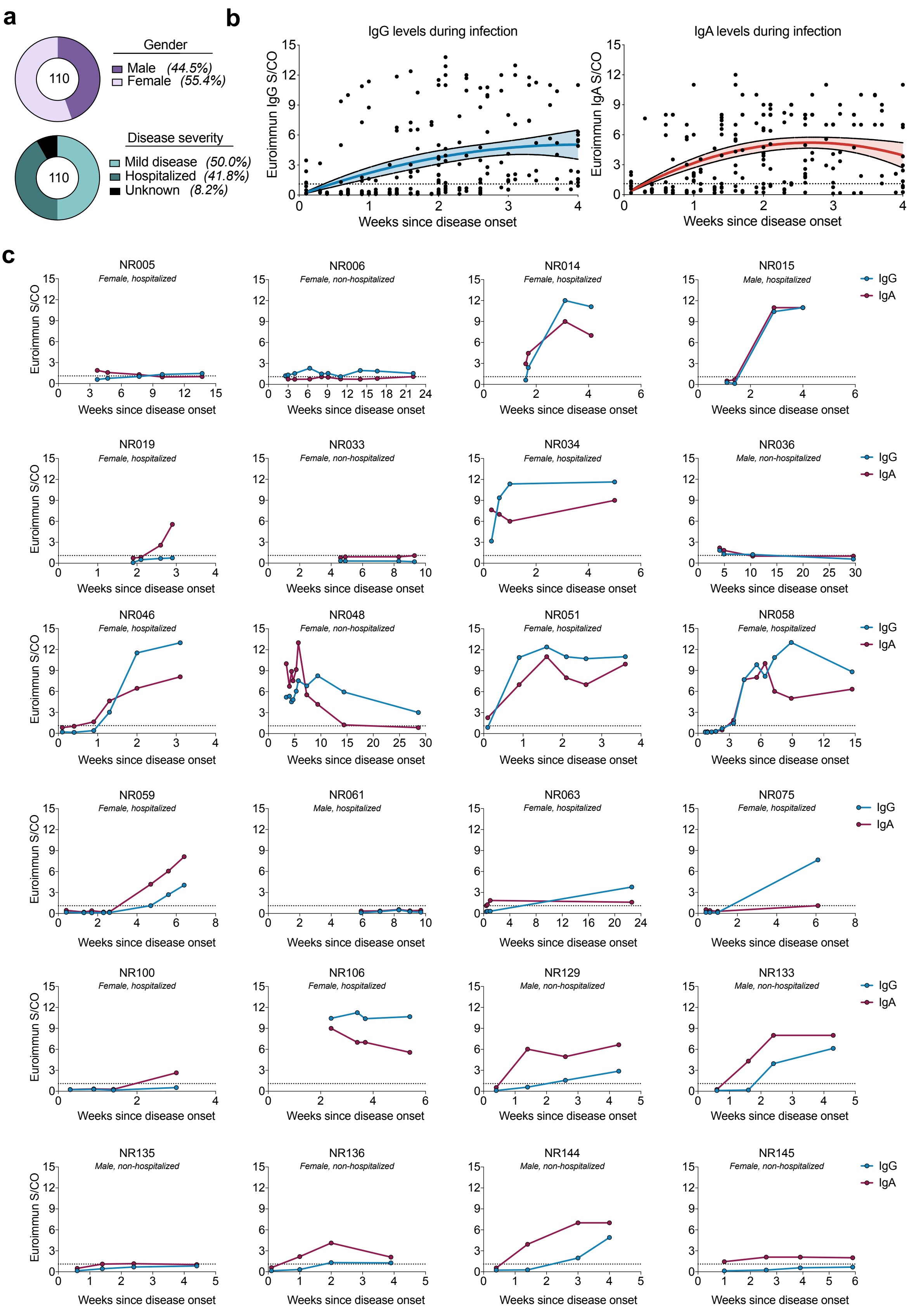


**Extended Data Figure 6: Antibody kinetics in the early phase of SARS-CoV-2 infection**

**a,** Pie charts indicating distribution of gender and disease severity in individuals who were longitudinally monitored starting from the early phase of infection. **b**, Plots depicting IgG and IgA ratios over time in individuals (n=107). Dotted line represents the limit of detection (S/CO=1.1) of the Euroimmun ELISA. Statistical analysis was performed using a second order polynomial quadratic equation (R2=0.128 for IgG and R2=0.140 for IgA) with 95% confidence interval shading (IgG in blue and IgA in red) of the best line. **c**, Individual plots depicting IgG (blue) and IgA (red) levels over time. Gender and disease severity are indicated within each plot. Dotted line represents the limit of detection (S/CO=1.1) of the Euroimmun ELISA.

### Extended data figure 7

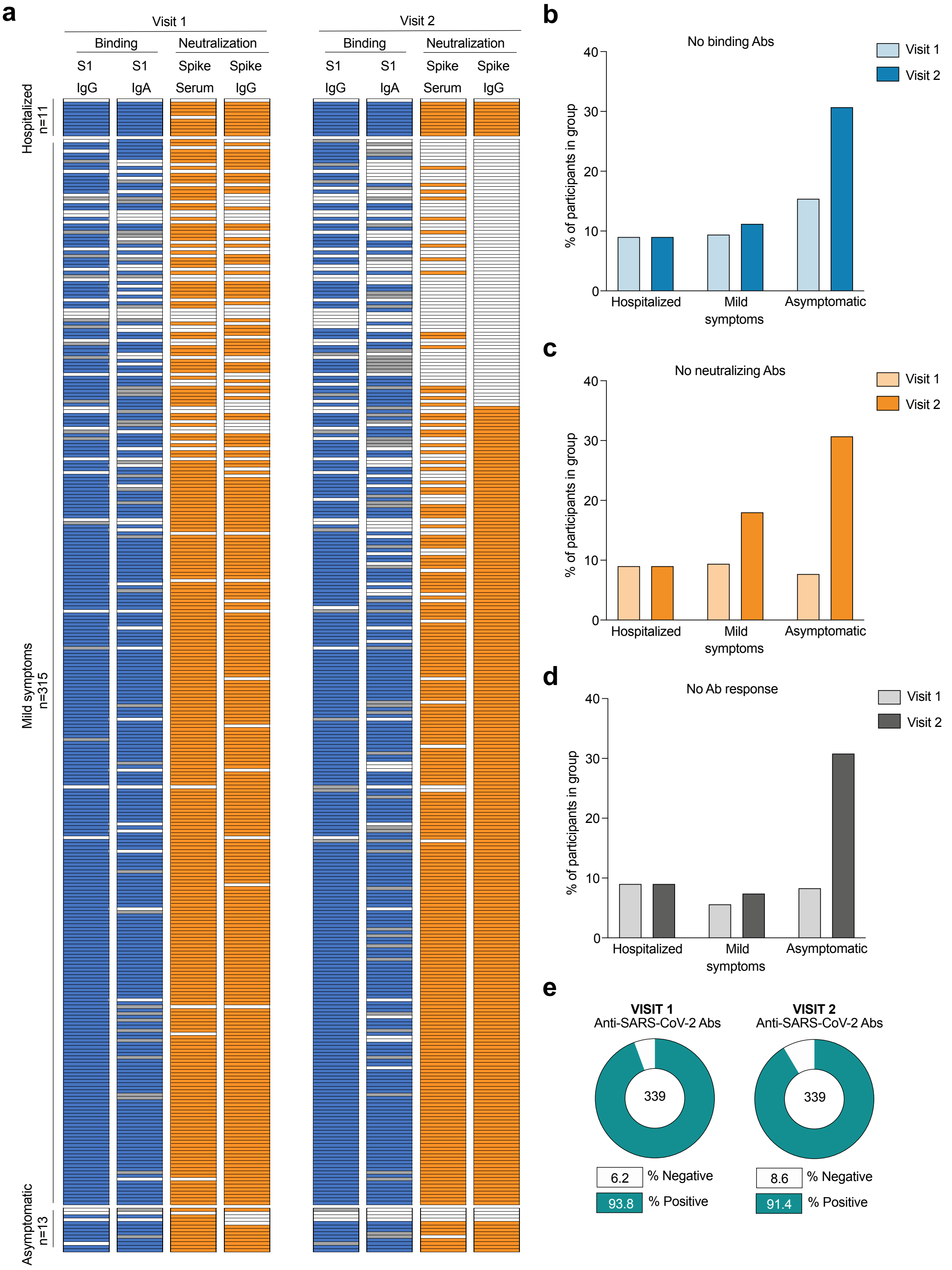


**Extended Data Figure 7: Changes in Ab response against SARS-CoV-2 over time**
**a,** plot depicting SARS-CoV-2 S1 binding and Wu01 pseudovirus neutralization of 339 individuals at visit 1 and visit 2 with each row representing 1 individual. Bar graphs showing change in fraction of individuals negative for anti-spike Abs (**b**), anti-spike NAbs (**c**) or any Ab response (**d**). **e**, pie chart evaluating the total presence if Ab response between visit 1 and 2 for all individuals.

#### Supplementary Table 1:Study cohort demographics, clinical information and antibody measurements

**Study cohort**

***Study cohort demographics*** ***n=963***

| Gender | | Age | |
|---|---|---|---|
| Male (n=) / (%) | Female (n=) / (%) | Median (years) | Range (years) |
| 444 / 46.1% | 519 / 53.9% | 44 | 18-79 |

***Study cohort clinical information***

| Asymptomatic (n=) / (%) | Symptomatic (n=) / (%) | Unknown (n=) / (%) |
|---|---|---|
| 44 / 4.5% | 911 / 94.6% | 8 / 0.8% |

| Fever (n=) / (%) | Cough (n=) / (%) | Sore throat (n=) / (%) | Rhinitis (n=) / (%) | Muscle ache (n=) / (%) | Headache (n=) / (%) | Diarrhea (n=) / (%) | Change in taste (n=) / (%) | Change in smell (n=) / (%) |
|---|---|---|---|---|---|---|---|---|
| 428 / 44.4% | 613 / 63.7% | 353 / 36.7% | 332 / 34.5% | 508 / 52.8% | 507 / 52.6% | 182 / 18.9% | 563 / 58.5% | 517 / 53.7% |

| Not hospitalized (n=) / (%) | Hospitalized (n=) / (%) | ICU (n=) / (%) | Unknown (n=) / (%) |
|---|---|---|---|
| 927 / 96.3% | 28 / 2.9% | 7 / 0.7% | 8 / 0.8% |

| Pre-conditions (n=) / (%) | No pre-conditions (n=) / (%) | Unknown (n=) / (%) |
|---|---|---|
| 224 / 23.3% | 695 / 72.2% | 44 / 4.6% |

| Respiratory (n=) / (%) | Heart (n=) / (%) | Kidney (n=) / (%) | Clotting (n=) / (%) | Cancer (n=) / (%) | Diabetes (n=) / (%) | Hypertension (n=) / (%) | Auto-immune (n=) / (%) | Other (n=) / (%) |
|---|---|---|---|---|---|---|---|---|
| 53 / 5.5% | 14 / 1.5% | 5 / 0.5% | 11 / 1.1% | 24 / 2.5% | 12 / 1.2% | 73 / 7.6% | 33 / 3.4% | 33 / 3.4 % |

**Study Visit 1**

***Neutralization analyse*** ***n=963***

| Time since disease onset | Serum/plasma ID50 | | | | Total IgG IC50 | | | |
|---|---|---|---|---|---|---|---|---|
| Median (weeks) | Geo. Mean (dilution) | Range (dilution) | Neutralizer (n=) | Non-neutralizer (n=) | Geo. Mean (ug/ml) | Range (ug/ml) | Neutralizer (n=) | Non-neutralizer (n=) |
| 7.3 | 111,3 | 10-90092 | 793 | 170 | 161 | 0.7-745.6 | 760 | 203 |

***ELISA analyses*** ***n=963***

| Time since disease onset | IgG | | | | IgA | | | |
|---|---|---|---|---|---|---|---|---|
| Median (weeks) | Geo. Mean (S/CO) | Pos (n=) | Neg (n=) | Equivocal (n=) | Geo. Mean (S/CO) | Pos (n=) | Neg (n=) | Equivocal (n=) |
| 7.3 | 4,1 | 797 | 122 | 44 | 3,0 | 676 | 189 | 98 |

**Study Visit 2**

***Neutralization analyse*** ***n=342***

| Time since disease onset | Serum/plasma ID50 | | | | Total IgG IC50 | | | |
|---|---|---|---|---|---|---|---|---|
| Median (weeks) | Geo. Mean (dilution) | Range (dilution) | Neutralizer (n=) | Non-neutralizer (n=) | Geo. Mean (ug/ml) | Range (ug/ml) | Neutralizer (n=) | Non-neutralizer (n=) |
| 17,3 | 64,3 | 10-5826 | 241 | 101 | 186,2 | 13.6-750 | 256 | 86 |

***ELISA analyses*** ***n=616***

| Time since disease onset | IgG | | | |
|---|---|---|---|---|
| Median (weeks) | Geo. Mean (S/CO) | Pos (n=) | Neg (n=) | Equivocal (n=) |
| 18,8 | 3,4 | 479 | 100 | 37 |

**Study Visit 3**

***ELISA analyses*** ***n=430***

| Time since disease onset | IgG | | | |
|---|---|---|---|---|
| Median (weeks) | Geo. Mean (S/CO) | Pos (n=) | Neg (n=) | Equivocal (n=) |
| 30,1 | 2,9 | 322 | 74 | 35 |

**Study Visit 4**

***ELISA analyses*** ***n=137***

| Time since disease onset | IgG | | | |
|---|---|---|---|---|
| Median (weeks) | Geo. Mean (S/CO) | Pos (n=) | Neg (n=) | Equivocal (n=) |
| 37,9 | 3,0 | 100 | 28 | 9 |
